# Upstream open reading frame inactivation augments GATA4 translation and cardiomyocyte hypertrophy in mice

**DOI:** 10.1101/2025.05.18.654700

**Authors:** Omar M. Hedaya, Feng Jiang, Uday Baliga, Aleksandr Ivanov, Si Chen, Jack L. Schwartz, Yui Kawakami, Chen Yan, Peng Yao

## Abstract

Upstream open reading frames (uORFs) are short peptide-encoding sequences located in the 5’ untranslated region (5’ UTR) of mRNAs, enabling translational repression of main (m)ORFs. While uORFs are found in ∼50% of mRNAs in humans, our understanding of their biological function remains limited. This study aims to elucidate the role of the uORF in the 5’ UTR of the Gata4 (GATA binding protein 4) gene in cardiac biology by inactivating its start codon (ΔuORF) in the mouse genome. Our investigation reveals that mice with Gata4 uORF inactivation manifest spontaneous cardiac hypertrophy without apparent fibrosis as they age. Utilizing single-nucleus RNA sequencing (snRNA-seq), we uncovered significant transcriptional variations between wild-type (WT) and ΔORF mice. Notably, mRNAs associated with sarcomeres and contractile functions show heightened expression levels, reflecting the hypertrophic phenotype. Notably, at least nine upregulated genes are GATA4-bound targets in mouse ventricles. Functional assessments of isolated primary adult cardiomyocytes confirmed enhanced hypertrophy and contractility in ΔORF mice. Additionally, we employed single-nucleus transposase-accessible chromatin (snATAC)-seq to investigate changes in chromatin accessibility. Our results indicated increased accessibility within specific transcription-regulatory elements linked to elevated gene transcription. These putative cis-regulatory elements (pCREs) are significantly enriched in MEF2 (myocyte enhancer factor 2) binding motifs. *In vitro* luciferase reporter assays further supported the regulatory potential of three of these pCREs, highlighting their role in the transcriptional enhancement of three GATA4 target genes bound by MEF2 and GATA4. These findings illuminate the role of uORF in negatively regulating GATA4 protein expression and cardiomyocyte hypertrophy at the organismal level and provide a novel therapeutic target for cardiac pathogenesis.

## Introduction

GATA binding protein 4 (GATA4), a zinc-finger transcription factor, is significantly involved in cardiac development, myocardial differentiation, cardiomyocyte (CM) hypertrophic growth and survival, and stress-responsive angiogenesis within the adult heart ^1–4^. Its significance is highlighted by its high expression levels in both embryonic and adult CMs, where it serves as a key transcriptional regulator for numerous cardiac genes, including atrial natriuretic factor (ANF), b-type natriuretic peptide (BNP), α-myosin heavy chain (α-MHC, also MYH6), β-myosin heavy chain (β-MHC, also MYH7), vascular endothelial growth factor A (VEGF-A), among others ^5,6^. GATA4 also plays a critical role in activating gene expression in response to pathological hypertrophic stimuli in CMs, including pressure overload and neurohumoral stimulation (e.g., isoproterenol, phenylephrine, and endothelin) ^7^. Studies utilizing the overexpression of GATA4 in cultured primary rat CMs and transgenic mice have shown its sufficiency in inducing cardiac hypertrophy ^6,8^.

Given its biological significance, GATA4 expression is subject to tight regulation. Various stimuli associated with cardiac hypertrophy and heart failure enhance GATA4 transcriptional activity through phosphorylation ^5^. For example, multiple hypertrophic stimuli trigger phosphorylation of GATA4 at serine 105 (Ser^105^), leading to enhanced DNA binding and transactivating activity ^9–12^. Ser^105^ phosphorylation of GATA4, facilitated by extracellular signal-regulated kinase (ERK) 1/2 and p38 mitogen-activated protein kinase (MAPK), exemplifies this regulation ^5,11^. Additionally, glycogen synthase kinase 3β negatively regulates GATA4 by reducing both basal and isoproterenol-induced nuclear expression of GATA4 by promoting its nuclear export ^13^. In mammalian cells, GATA4 is regulated through the autophagy-lysosome pathway, where p62, an autophagy adaptor, is responsible for the degradation of GATA4 ^14^. In certain cellular events such as aging, ATM and ATR inhibit this mechanism, stabilizing GATA4 and activating a senescence gene program ^14^.

GATA4 also shows regulation at the level of translation initiation. Translation initiation is the process by which the small 40S ribosomal subunit is loaded onto the mRNA at the 5’ cap, scans along the 5’-UTR until reaching a start codon, and associates with the large 60S ribosomal subunit to create the complete 80S ribosome required for productive translation. Previously, we have shown that GATA4 contains an upstream open reading frame (uORF) in its 5’-UTR which regulates its translation^1^. uORFs are peptides encoded from a start codon within the 5’-UTR of protein-coding mRNAs. Translation of these uORFs is typically repressive to translation of the protein-coding main (m)ORF. In the prevailing model of uORF-mediated translational repression, uORFs are repressive by providing an alternative site which the 40S ribosome can associate with the 60S ribosome and initiate productive translation. Once ribosomes have finished translating the uORF peptide, they dissociate from the mRNA and never reach the mORF to translate the functional protein. In this model, mORF translation of uORF-containing mRNAs is possible due to leaky scanning of the 40S ribosome through uORF start codons, allowing for initiation at the mORF rather than the uORF, or through re-initiation after translation of the uORF peptide ^2–6^.

Previous work has confirmed that the GATA4 uORF is translated into a small peptide in both human and mouse hearts ^15,16^. However, the GATA4 uORF-encoded peptides are not conserved between humans (9 amino acids) and mice (18 amino acids), implying a critical regulatory function driven by the translation event of this uORF ^17^ rather than a specific bioactivity of the uORF-encoded peptide ^18^. Our previous studies revealed that the 11-nt sequence downstream of the uORF AUG start codon is highly conserved and forms double-stranded (ds)RNA to promote translation of the uORF ^17^. The human recombinant uORF-derived peptide has been shown to preferentially bind to nuclear proteins, including multiple RNA-binding proteins (e.g., ADAR, RBM14, UBAP2L, XRN1), suggesting its potential nuclear function in the CM RNA metabolism; however, it requires further validation ^18^.

Exploring the impact of uORFs on biological functions is a complex endeavor, largely limited to *in vitro* studies ^18–22^. In this study, we investigated the effects of inactivating the *Gata4* uORF in a genetic mouse model, employing CRISPR-Cas9 technology by modifying the first nucleotide of the AUG codon from ‘A’ to ‘U’ to inactivate this start codon. We demonstrated that this uORF intrinsically acts as an anti-hypertrophic element and reduces the endogenous synthesis of the GATA4 protein. Through echocardiographic and histological evaluations, we observed a gradual development of cardiac hypertrophy without notable fibrosis in these uORF loss-of-function mice over a 12-month period, which becomes more pathological under increased cardiac pressure overload stress induced by transverse aortic constriction surgery. Utilizing single-nucleus RNA sequencing (snRNA-seq) and the Assay for Transposase-Accessible Chromatin (snATAC-seq), we identified shifts in gene expression profiles, specifically in left ventricular CMs, suggesting a more hypertrophic and contractile phenotype in this mouse model at baseline. The snATAC-seq data allows us to pinpoint changes in chromatin accessibility that correspond with alterations in mRNA levels identified by snRNA-seq. Our results imply that the uORF modulates GATA4 protein levels and consequently affects chromatin accessibility of potential cis-regulatory elements recognized by GATA4 or its co-factors MEF2A and MEF2C, thereby enhancing transcriptional activity and modulating the expression of a specific cohort of genes. Validation of these sequences upstream of a minimal SV40 promoter through a luciferase reporter assay further substantiates their role in amplifying transcription. This study underscores the importance of GATA4 uORF in translational control and gene regulation in the heart.

## Results

### Generation of *Gata4* ΔuORF knock-in mice with enhanced mORF translation

Both human and mouse GATA4 5’ UTRs contain a uORF with an in-frame start codon ATG and stop codon TAA (in humans) or TGA (in mice) with no conserved peptide-coding sequences in between (**Figure 1A**). To evaluate whether the uORF in the mouse *Gata4* 5’ UTR exerts a comparable inhibitory influence on mRNA translation as observed in human systems, such as human AC16 cardiomyocyte cells and embryonic stem cell-derived cardiomyocytes ^17^, we conducted experiments using reporter plasmids containing either the wild-type (WT) mouse *Gata4* 5’ UTR or a version lacking the uORF (ΔuORF) (**Figure 1B**). Our findings showed a significant 2.6-fold increase in luciferase reporter activity, while mRNA levels remained unchanged (**Figure 1C**). This confirms the suppressive capability of the mouse Gata4 uORF in a manner consistent with observations in human cells ^17^.

**Figure 1.**
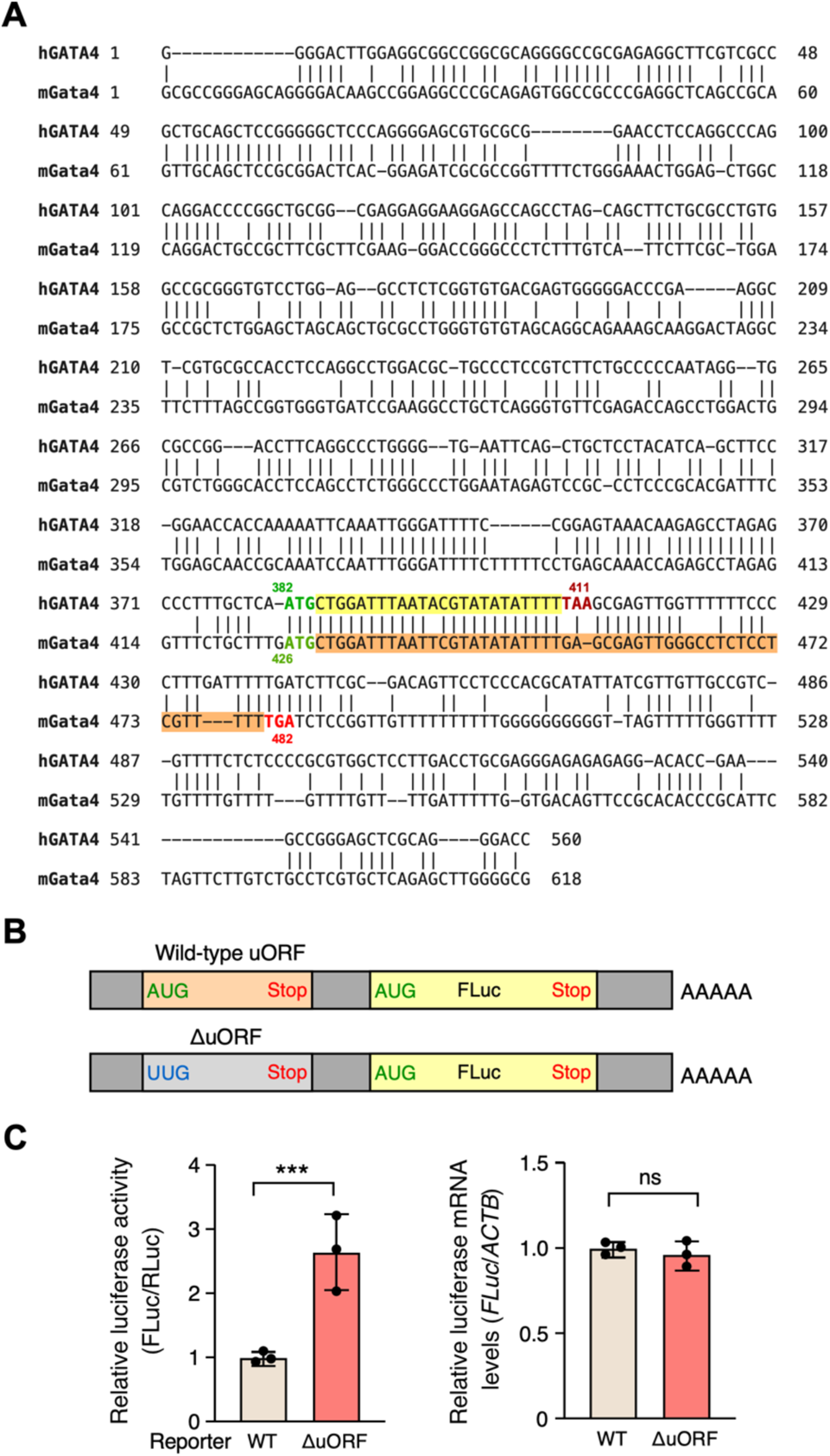
Mouse Gata4 uORF serves as a repressive regulatory element for mORF translation. **A.** Pair-wise alignment of human and mouse GATA4 5’-UTR sequences. The ATG start codon is highlighted in green, and the stop codons are red. The upstream (u)ORF after ATG is highlighted in yellow in human GATA4 and orange for mouse Gata4. **B**. Schematic depicting firefly luciferase (FLuc) reporter constructs that include the full-length 5’ UTR RNA sequence of mouse *Gata4* mRNA in both wild-type (WT) and ΔuORF (ATG-to-TTG start codon mutation) configurations. **C**. Left: Dual-luciferase reporter assay conducted in HEK293T cells, showing firefly luciferase (FLuc) activity in both WT and ΔuORF constructs, normalized to a Renilla luciferase (RLuc) control plasmid and calculated relative to the WT. Right: RT-qPCR results for FLuc mRNA, normalized to *ACTB*. Data are represented as mean ± SD. *** *P* < 0.001; Statistical significance was confirmed by an unpaired two-tailed Student *t* test for (N=3 biological replicates).

After recognizing the evolutionary conservation of this uORF in humans and mice, we introduced an identical ΔuORF mutation (**Figure 1B**) into a genetic mouse model. Utilizing CRISPR-Cas9 technology and a homology-directed repair DNA template featuring the ΔuORF sequence (**Figure 2A**), we also incorporated a mutation in the protospacer adjacent motif (PAM) to inhibit repetitive Cas9 cleavage post-editing. Following four breeding cycles to secure genetic uniformity, we successfully generated the targeted ΔuORF mouse model as indicated by PCR-derived amplification of the DNA fragment spanning the editing sites, followed by Sanger sequencing (**Figures 2B and S1A**). Within a group of 55 mice, we observed an expected Mendelian ratio of approximately 1: 2: 1 for WT, heterozygous, and homozygous knock-in (KI) mice (**Figure S1B**) derived from heterozygous parental mice, indicating that the mutation likely has no adverse effects on embryonic development and organismal viability. Western blotting performed on heart lysates from 8-week-old KI mice showed a dose-dependent increase in protein levels, contingent upon the absence of uORF copies in the genome, either heterozygous or homozygous, resulting in 1.6- and 2.3-fold increases in protein levels, respectively (**Figure 2B, C, D**, left panel). Similar to the luciferase reporter assay (**Figure 1A**), endogenous *Gata4* mRNA levels remained unaltered (**Figure 2D**, right panel), supporting a post-transcriptional regulatory mechanism consistent with uORF-mediated translational control.

**Figure 2.**
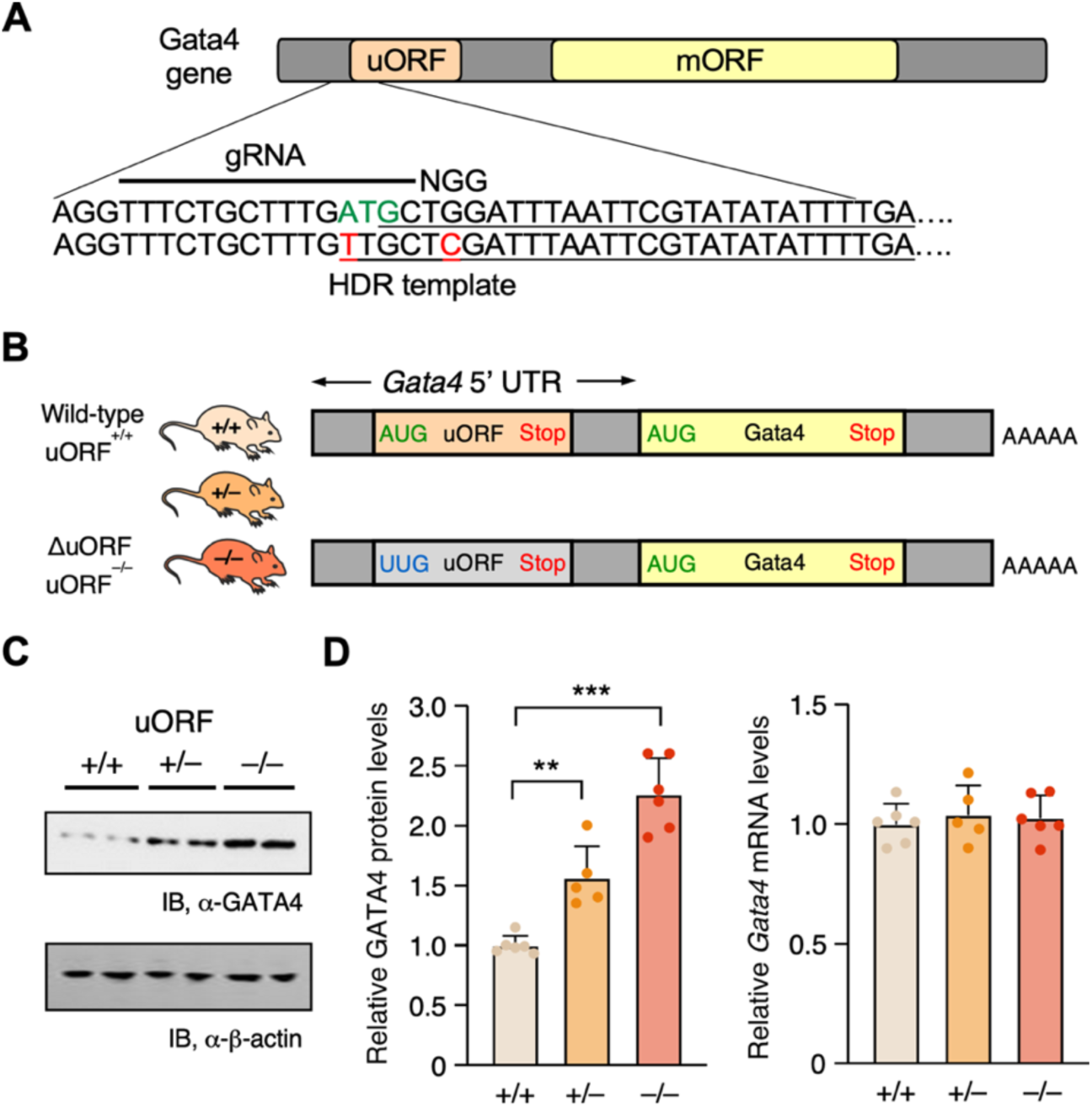
Inactivation of uORF elevates GATA4 protein levels in a knock-in mouse model. **A.** Schematic demonstrating the use of CRISPR-Cas9 technology coupled with homology-directed repair (HDR) for genomic inactivation of the *Gata4* uORF, featuring an ATG-to-TTG ΔuORF mutation and including the protospacer adjacent motif (PAM), using HDR DNA template, guide (g)RNA, with PAM. **B**. Schematic illustrating mRNA structure in mice following CRISPR-Cas9-mediated genomic inactivation of *Gata4* uORF via an ATG-to-TTG ΔuORF knock-in mutation. **C**. Western blot analysis of heart lysates, quantifying GATA4 protein levels in wild-type (+/+), heterozygous (+/-), and homozygous (-/-) mice with ΔuORF mutations; data are quantified in panel **D** (left). Corresponding *Gata4* mRNA levels, normalized to *Actb*, show no significant changes (right). Data are represented as mean ± SD. ** *P* < 0.01, *** *P <* 0.001; Statistical significance was confirmed by 2-way ANOVA followed by Holm-Sidak post hoc test (N = 6 mice; 3 males and 3 females).

### ΔuORF mice exhibit cardiac hypertrophy without any discernible cardiac fibrosis

GATA4 is primarily expressed in cardiomyocytes and germ cells according to the gene expression profiles across different cell types in the Human Protein Atlas database ^23^. To examine the cardiac phenotypes in adult ΔuORF KI mice, we undertook a year-long study starting with 4-week-old mice, utilizing echocardiography for longitudinal analysis (**Figure 3A**). Echocardiographic assessments were performed monthly for the first six months and bi-monthly for the following half-year. Intriguingly, we observed a gradual increase in left ventricular mass in homozygous ΔuORF KI mice, reaching up to 106.2 ± 6.69 mg, compared to 80.5 ± 23.69 mg in heterozygous KI mice and 69.6 ± 6.55 mg in WT mice at the end point of our observations (at the age of 12 months) (**Figure 3B**, left panel). Simultaneously, with a stable heart rate, we noted a rise in stroke volume between the second and fifth months, suggesting an increased cardiac output of 24.0 ± 3.95 ml/min in homozygous ΔuORF mice compared to 22.0 ± 5.90 ml/min in heterozygous (not significant) and 20.8 ± 2.93 ml/min in WT counterparts (at the age of 2 months). This finding may imply enhanced cardiac contractility (**Figure 3B**, middle panel). Intriguingly, the significance of the cardiac output difference among the three genotypes did not hold after month 5. Notably, ejection fraction metrics remained stable, indicating cardiac hypertrophy without a decline in cardiac contractile function or subsequent heart failure (**Figure 3B**, right panel).

**Figure 3.**
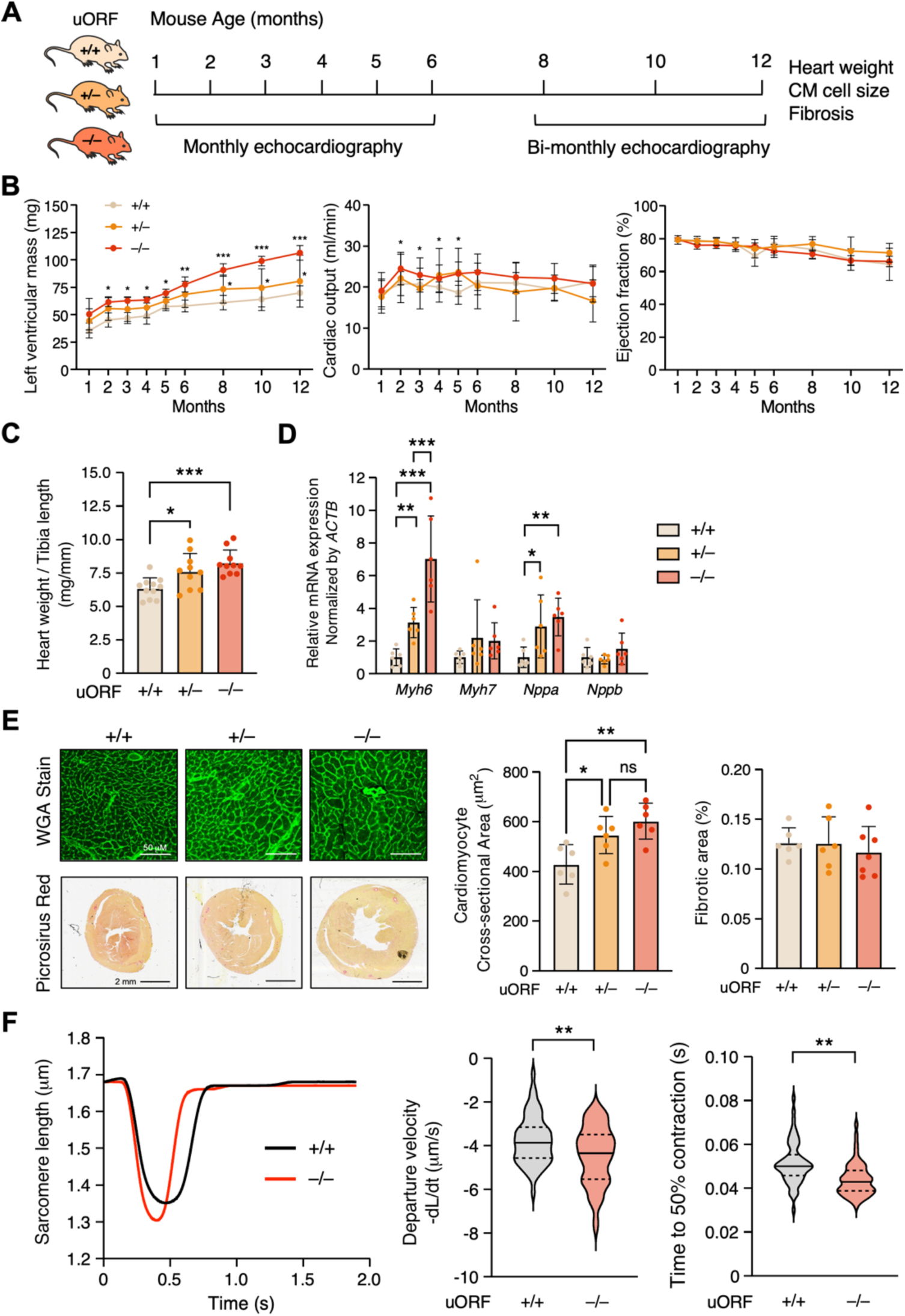
*Gata4* ΔuORF mice develop spontaneous cardiac hypertrophy without fibrosis. **A.** Schematic illustrating the frequency of echocardiographic measurement of 4-week-old wild-type (WT; +/+), heterozygous (+/-), and homozygous (-/-) mice over 12 months. **B**. Measurement of left ventricular mass, cardiac output, and ejection fraction in three groups of mice. **C-E**. Experiments were conducted on harvested hearts at the 12-month endpoint. **C**. Heart weight normalized to the tibia length as a gross measurement of cardiac hypertrophy. **D**. RT-qPCR of the hypertrophy markers: *Myh6, Myh7, Nppa,* and *Nppb* mRNA levels normalized to *Actb*. **E**. Left upper panel: Immunomicrograph of mouse cross-sections stained with wheat germ agglutinin (WGA)-Alexa fluor 480 to highlight cellular cross-sectional area quantified in the middle panel. Left lower panel: A scan of heart tissue slides was stained with picrosirius red, highlighting collagen deposition and quantified in the right panel. **F**. *Gata4* uORF inactivation enhanced the contractility of isolated CMs at the baseline as indicated by Sarcomere length relative to time, Departure velocity, and Time to 50% contraction. N = ∼100 CMs were quantified for WT and homozygous (-/-) mice from 3 hearts. Data are represented as mean ± SD. ** *P* < 0.05, ** *P* < 0.01, *** *P <* 0.001; Statistical significance was confirmed by 2-way ANOVA followed by Holm-Sidak post hoc test. N = 11 (6 females and 5 males), N = 10 (6 females and 4 males), and N = 11 (5 females and 6 males) for WT, heterozygous, and homozygous mice, respectively, in **B**. N = 6 (3 males and 3 females) in **C-E**.

At the one-year time point, consistent with Figure 3B (left panel), we observed a dose-dependent increase in heart weight relative to tibia length (**Figure 3C**): 8.3 ± 0.96 mg/mm for homozygous mice, 7.6 ± 1.34 mg/mm for heterozygous mice, and 6.3 ± 0.96 mg/mm for WT mice. We detected an increase in biomarker mRNA expression indicative of cardiac hypertrophy, specifically *Myh6* and *Nppa*, while the levels of *Myh7* and *Nppb* remained unchanged (**Figure 3D**). Furthermore, using wheat germ agglutinin (WGA) staining to delineate cellular boundaries and quantification through ImageJ, we found an enlargement in the cross-sectional area of cardiomyocytes from 427.7 ± 79.34 μm^2^ (WT mice) to 545.4 ± 74.61 μm^2^ and 601.0 ± 72.33 μm^2^ in heterozygous and homozygous ΔuORF mice, respectively (**Figure 3E**, left and middle panels). In contrast, picrosirius red staining, which emphasizes collagen deposition, showed no statistically significant rise in myocardial regions as quantified through ImageJ (**Figure 3E**, left and right panels). In agreement with upregulated GATA4 protein, homozygous ΔuORF mouse CMs exhibited enhanced contractile function compared to WT CMs at baseline (**Figure 3F**). The time for one contraction-relaxation cycle is shorter in ΔuORF mouse CMs than that in WT CMs (**Figure 3F**, left panel). Accordingly, the departure velocity and time to 50% contraction shortening were decreased in ΔuORF CMs (**Figure 3F**, middle and right panels). Taken together, these results suggest that genetic inactivation of *Gata4* uORF in mice leads to upregulated expression of GATA4 protein, augmented cardiac hypertrophy, and enhanced contractile function in mouse CMs.

### *Gata4* ΔuORF mice exhibit decompensatory responses to cardiac pressure overload

To further elucidate the role of *Gata4* uORF in stress-induced cardiac hypertrophy, we subjected mice from three genotype groups to transverse aortic constriction (TAC). Compared to WT mice, levels of GATA4 protein were significantly elevated in the hearts of heterozygous and homozygous ΔuORF mice with pressure overload stimulations (**Figure S2A**). In contrast, mRNA levels remained comparable among the three groups (**Figure S2B**).

We monitored cardiac function at baseline and biweekly over 8 weeks after the TAC surgery using echocardiography (**Figure S2C-E**). At baseline, the groups showed no significant differences in ejection fraction and left ventricular mass (**Figure S2C, D**). However, elevated cardiac output was observed in homozygous ΔuORF mice at baseline (**Figure S2E**), corroborating our previous findings in Figure 3C. A distinct shift became evident following the onset of pressure overload (**Figure S2C-E**). In particular, there was a pronounced increase in cardiac hypertrophy alongside a steep decline in ejection fraction, most notably in homozygous ΔuORF mice. Specifically, post-TAC ejection fractions at 8 weeks were 41.4% ± 13.84 in homozygotes, 65.1% ± 13.49 in heterozygotes, and 58.5% ± 13.78 in WT mice (**Figure S2C**). Similarly, left ventricular mass increased significantly post-TAC at 8 weeks, with values of 183.8 mg ± 33.87 in homozygotes, 106.4 ± 23.19 mg in heterozygotes, and 115.0 ± 35.73 mg in WT mice (**Figure S2D).** Interestingly, the decline in cardiac output was comparable across all three groups (**Figure S2E**).

Upon evaluating heart weight normalized to tibia length, we found it stable at baseline across all groups, with a slight increase in the homozygous ΔuORF mice (**Figure S2F**). However, at 8 weeks post-TAC, a significant increase in heart weight was exclusively observed in the homozygous ΔuORF mice, highlighting the crucial role of GATA4 uORF in limiting cardiac hypertrophy (**Figure S2F**). Consistent with this, WGA staining revealed a marked enlargement in the cardiomyocyte cross-sectional area, indicative of hypertrophic growth, while picrosirius red staining showed a significant increase in interstitial fibrosis (**Figure S2G-I**). These observations collectively suggest that ΔuORF mice hearts are more susceptible to hypertrophic remodeling under pressure overload. This heightened vulnerability may cause these hearts to transition more rapidly from compensatory to decompensatory stages during cardiac hypertrophy, thereby accelerating the progression to heart failure.

### Single nucleus analysis of transcriptome and chromatin accessibility in adult WT and ΔuORF KI mouse hearts

As a transcription factor prevalent in the heart, GATA4 is intricately linked with chromatin remodeling and transcriptional alterations affecting various cardiac cells ^24^. Consequently, we performed an in-depth examination of chromatin accessibility and gene transcription changes across distinct cardiac cell types in WT and homozygous ΔuORF KI mice. Our approach utilized simultaneous single-nucleus ATAC-sequencing (snATAC-seq) and single-nucleus RNA-sequencing (snRNA-seq) within each nucleus, known as single-nucleus dual-omics analysis (**Figure 4A, Table S1**). This method enabled us to assign the same identifier to RNA-seq reads and ATAC-seq fragments originating from identical nuclei, allowing precise matching of gene expression with chromatin accessibility. We gathered expression and accessibility profiles for 6,473 nuclei, with a median of 837 RNA-seq reads and 1,343 ATAC-seq fragments per nucleus. These nuclei comprise 3,373 nuclei from WT hearts and 3,100 nuclei from homozygous ΔuORF KI hearts.

**Figure 4.**
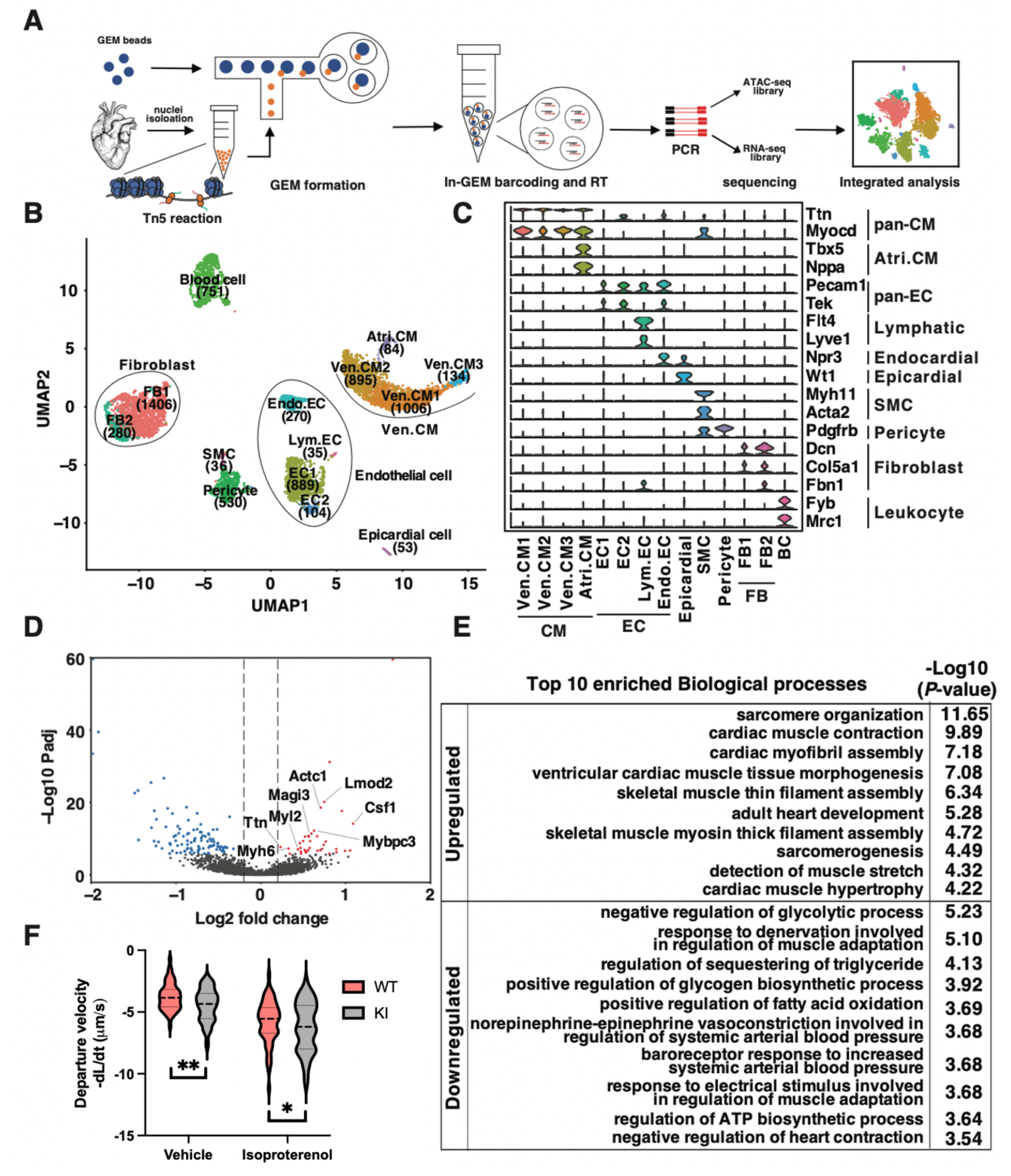
snRNA/ATAC-seq analysis of wildtype and ΔuORF mouse hearts. **A.** A sketch showing the workflow of snRNA/ATAC-seq. **B**. A UMAP presentation of clustering results based on the top 3000 variant features. Fourteen clusters were identified and labelled according to cell type and abundance. **C**. Violin Plots showing the expression of well-established marker genes in each cluster. Scaled and normalized expressions were plotted on the y-axis. **D**. A volcano plot showing the differentially expressed genes (DEGs) in Ven.CM. Genes with significantly increased expression in ΔuORF KI hearts (Log_2_ fold change > 0.2 and Bonferroni correction adjusted *P*-value < 0.05) were red-colored. Genes with significantly decreased expression (Log_2_ fold change < −0.2 and Bonferroni correction adjusted *P*-value < 0.05) in ΔuORF hearts were colored blue. This is the zoom-in version of the same plot shown in Figure S3F. Published GATA4 targets are highlighted in the volcano plot. **E**. Gene ontology analysis results showing the top 10 enriched biological processes in upregulated and downregulated DEGs. **F**. violin plots show the contraction parameters of ∼100 cardiomyocytes from 3 mice at the baseline (Figure 3F, middle panel) and upon isoproterenol (ISO; 10 μM) stimulation. The contraction velocity of WT versus ΔuORF KI cardiomyocytes was calculated as the rate of change of cell size over time. Data in this figure are represented as mean ± SD. * *P* < 0.05, ** *P* < 0.01; Statistical significance was confirmed by 2-way ANOVA followed by Holm-Sidak post hoc test. Ven.CM, ventricular cardiomyocytes; Atri.CM, atrial cardiomyocytes; FB, fibroblasts; EC, endothelial cells; SMC, smooth muscle cells; BC, blood cells.

By utilizing Signac and Seurat, we identified 14 distinct clusters, which we subsequently annotated based on the RNA expression patterns of established lineage-specific marker genes (**Figures 4B and 4C, Table S2**). This annotation was further supported by gene accessibility data summarizing chromatin accessibility of the promoter and gene body of these markers (**Figure S3A**). Notably, we observed a high correlation between gene accessibility and expression within the same or closely related clusters, underscoring the accuracy of our clustering and annotation approach (**Figure S3B**). These 14 clusters collectively represent seven major cardiac cell types: ventricular cardiomyocytes (Ven.CM), atrial cardiomyocytes (Atri.CM), fibroblasts (FB), endothelial cells (EC), smooth muscle cells (SMC), pericytes, and epicardial cells. Importantly, the abundances of nuclei in these cell types did not show significant changes in ΔuORF hearts compared to WT hearts, except for endothelial and fibroblast cells (**Figure S3C and S3D**).

Subsequently, we examined the differential gene expression patterns between WT and ΔuORF KI hearts. While GATA4 is expressed in both atrial and ventricular cardiomyocytes (**Figure S3E**), the majority of gene expression differences were observed in Ven.CM (**Figure S3F, Table S3**). We identified 40 upregulated and 100 downregulated differentially expressed genes (DEGs) in ΔuORF Ven.CM, with |Log2 fold change| > 0.2 and Bonferroni correction adjusted *P*-value < 0.05 (**Figure 4D, Table S3**). After overlapping the 40 upregulated genes in ΔuORF Ven.CM with a well-established ChIP-seq-derived GATA4 target genes in mouse ventricles (1801 genes), we identified nine GATA4 target genes, including Ttn, Myh6, Myl2, Mybpc3, Actc1, Lomd2, Magi3, Ankrd1, and Csf1 ^25^, most of which are classical cardiomyocyte sarcomere genes required for contractile functions. The upregulated DEGs include well-established GATA4 targets, reflecting the role of GATA4 as a transcription factor (**Figure 4D**). Our gene ontology (GO) analysis suggested that the upregulated DEGs were enriched in biological processes related to cardiac muscle contraction and hypertrophy. In contrast, downregulated DEGs were associated with processes related to the negative regulation of glycolysis, muscle adaptation regulation, and negative regulation of heart contraction, considered as potential secondary or compensatory effects (**Figure 4E**). Consistent with the changes in gene expression profiles, contractility was significantly enhanced in KI heart-derived primary cardiomyocytes compared to control WT cells (**Figure S3G**), indicating a functional impact of uORF activity on regulating GATA4 protein expression and cardiomyocyte contractile activity.

### Identification of cis-regulatory DNA elements with altered accessibility in ΔuORF ventricular cardiomyocytes

To identify accessible chromatin regions, we aggregated the snATAC-seq fragments from individual clusters and employed MACS2 for peak calling (**Figure 4A, Table S4**). Our analysis revealed a total of 72,219 universally accessible chromatin regions and 7,155 regions specific to cardiomyocytes (CM), fibroblasts (FB), endothelial cells (EC), pericytes, and blood cells (BC) (**Figure S4A**). Subsequent gene ontology analysis indicated that these cell type-specific regions were proximal to genes involved in relevant biological processes (**Figure S4B**). We then searched for putative cis-regulatory elements (pCREs) within the regions by calculating the correlation between each chromatin region and the mRNA expression of nearby genes (**Figure S4C, Table S5**). Among all 79,374 regions, we identified 7,264 region-gene correlated pairs, encompassing 6,391 regions and 3,630 genes. These 6,391 pCREs exhibited significant positive correlations with the expression of their paired genes, strongly indicating their potential role as enhancers for the correlated genes.

We subsequently investigated changes in pCRE accessibility in ΔuORF KI Ven.CM. Among the 6,391 pCREs analyzed, 193 exhibited significantly altered accessibility (*P* < 0.05) in ΔuORF KI Ven.CM compared to the WT control CM (**Table S6**). This subset comprised 58 pCREs with increased accessibility and 135 with decreased accessibility (**Figure 5A**). A strong positive correlation between mRNA expression and chromatin accessibility of eight exemplary genes was highlighted, out of which seven were GATA4 target genes except Pcbpn1 (**Figure 5A**, lower panel). Notably, 33 of the 193 pCREs were correlated with DEGs in Ven.CM. Among these, 31 exhibited coordinated changes in ΔuORF KI hearts, where both the pCRE and its correlated DEG either increased or decreased, while the remaining pCREs showed opposite changes (**Figure S4D**).

**Figure 5.**
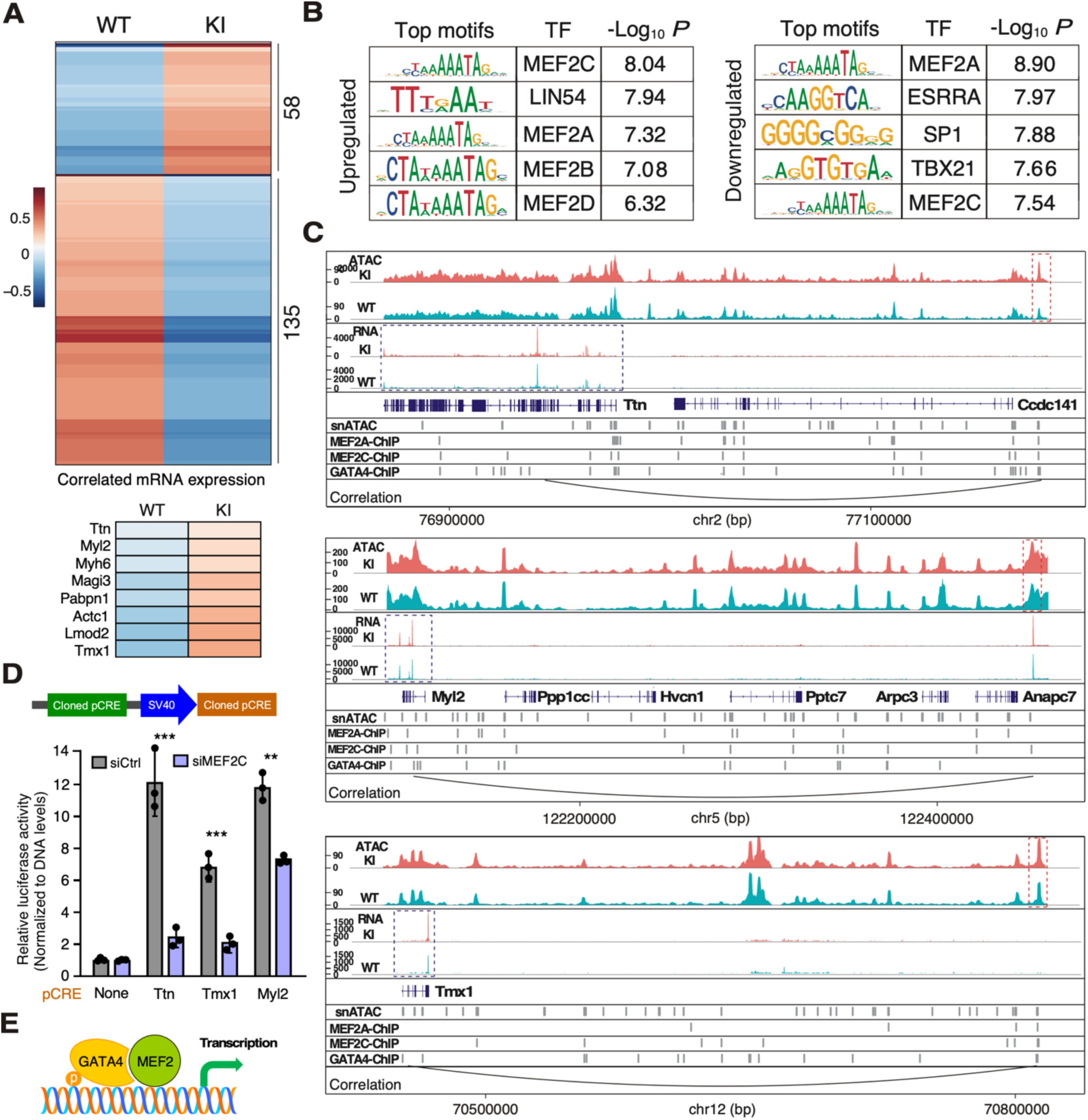
Identification of cis-regulatory DNA elements with altered accessibility in ΔuORF ventricular cardiomyocytes. **A.** A heatmap showing the accessibility of putative cis-regulatory elements (pCREs) with altered accessibility in ΔuORF Ven.CM. Scaled and normalized accessibility were plotted. Correlated mRNA expression with chromatin accessibility of multiple GATA4 target genes was highlighted. **B**. Motif analysis results showing the top 5 enriched transcription factor binding motifs in upregulated (left) and downregulated (right) pCREs. JASPAR2023 was used as the reference dataset for transcription factor binding motifs. **C**. Snapshots of the mouse genome showing examples of a pCRE and its correlated gene. Each snapshot includes the coverage of snATAC-seq fragments (1^st^ and 2^nd^ row), the coverage of snRNA-seq reads (3^rd^ and 4^th^ row), gene annotation (5^th^ row), accessible regions identified in snATAC-seq (6^th^ row), binding sites of MEF2A, MEF2C, and GATA4 identified by published ChIP-seq (7^th^ to 9^th^ row), and a curve connecting the pCRE with correlated gene. The red box indicates the pCRE, while the blue box indicates the correlated gene of the pCRE. For wildtype and ΔuORF, the snRNA-seq reads and snATAC-seq fragments from all nuclei in Ven.CM were summarized together for plotting. Normalized counts were plotted as the Y-axis for snATAC-seq, while RPKM values were plotted as the Y-axis for snRNA-seq. **D**. Firefly luciferase (FLuc) luminescence in the various reporters normalized to an empty vector containing the SV40 promoter. The values of each group were normalized to the DNA levels of FLuc. Upper panel: Schematic highlighting the reporter designed to test the identified pCREs. The pCREs were cloned upstream of a minimal SV40 promoter that drives a FLuc ORF. This data is represented as mean ± SD. * *P* < 0.05, ** *P* < 0.01; Statistical significance was confirmed by 2-way ANOVA followed by Holm-Sidak post hoc test. **E**. Schematic model of MEF2C-GATA4 complex in binding adjacent DNA regions for activating gene transcription.

Interestingly, both the increased and decreased pCREs showed a marked enrichment of MEF2A and MEF2C binding motifs (**Figure 5B**), strongly suggesting that these motifs serve as targets for the transcription factors MEF2A and MEF2C, whose expression remains unchanged in ΔuORF hearts (**Figure S4E**). This idea gains further support from the observation that a substantial portion of these pCREs also overlap with MEF2A and MEF2C binding targets identified through ChIP-seq (**Figure S4F, Table S7**) ^26^. Notably, many of these pCREs are also inferred to be GATA4 binding targets based on ChIP-seq data ^26^, even though the GATA4 binding motif is not notably enriched (**Figure S4F**).

To validate our identified pCREs, we cloned the pCREs that regulate Ttn, Myl2, and Tmx1 as enhancers into a luciferase reporter plasmid (**Figure 5D**, upper panel). Using the same amount of DNA, plasmids containing pCREs showed significantly increased luciferase expression (**Figure 5D**, lower panel). We performed the same experiment following the knockdown of MEF2A or MEF2C to examine their response to these factors. We noted significantly reduced luciferase reporter activity after the knockdown of MEF2C, suggesting a direct role for this transcription factor in regulating GATA4-dependent gene expression as a co-factor (**Figure 5E**) ^27,28^.

## Discussion

GATA4 is a transcription factor that plays an essential role in cardiac development; GATA4 knockout embryos exhibit defective heart tube morphogenesis, leading to embryonic arrest by day E9.0 ^29^. During heart development, GATA4 promotes the cardiac gene program while repressing others ^5,6,30^. In the adult heart, GATA4 regulates cardiac hypertrophy and the cellular response to stressors ^5–7,10^. On the other hand, GATA4 expression is also tightly regulated through various mechanisms at the post-translational level, including through post-translational modification and protein degradation ^5,10,11,13,14^. Our study investigated the previously unexamined regulatory role of the uORF in the 5’ UTR of the *Gata4* gene. We demonstrated that this uORF acts as a repressor of GATA4 protein expression and significantly influences the cardiac hypertrophic response mediated by GATA4. Using a CRISPR-generated mouse model with the *Gata4* uORF inactivated via a single-nucleotide mutation of the uORF start codon (from ATG to TTG), we observed increased levels of GATA4 protein without alteration of *Gata4* mRNA. The increased GATA4 protein expression triggered a persistent hypertrophic response throughout a 12-month study period. Furthermore, this elevated expression of *Gata4* heightened the heart’s susceptibility to hypertrophy under pressure overload, ultimately leading to cardiac decompensation and fibrosis. The regulatory dynamics of this uORF during various cellular events remain an open question. Previous research suggests that uORFs can be stress-responsive and regulated by RNA helicases, indicating potential contextual variance in their activity ^19,20,30,31^.

To understand the spontaneous hypertrophy observed in our mouse model at the molecular level, we utilized a single-nucleus dual-omics approach, which included snRNA-seq and snATAC-seq. Our analyses revealed that differentially expressed genes (DEGs) were primarily identified in ventricular myocytes, implying a ventricular-specific role for the GATA4 uORF. In contrast, the absence of DEGs in atrial myocytes may be due to limited sequencing coverage, a common limitation encountered in single-cell and single-nucleus sequencing technologies. Further unbiased methods are needed to comprehensively evaluate atrial myocyte gene expression. Several key myocyte structural and functional genes, including *Ttn*, *Actc1*, *Myl2*, and *Mybpc3*, fall under the regulatory purview of the *Gata4* mRNA uORF, and their increased expression in the ΔuORF mouse model likely contributes to enhanced myocyte hypertrophic growth and contractility. Interestingly, we also observed alterations in nuclear abundance in endothelial and cardiac fibroblast cells, despite their modest expression levels of *Gata4* mRNA, suggesting possible crosstalk among cell types, especially between cardiomyocytes and other cardiac cell types.

Although the exact molecular mechanism remains elusive, our snATAC-seq data suggest that the *Gata4* uORF may influence gene expression by modulating enhancer accessibility. We identified novel enhancers with altered accessibility following *Gata4* uORF inactivation, which regulates the transcription of key myocyte genes as established GATA4 targets. Interestingly, most of these enhancers function as binding sites for either MEF2A or MEF2C, transcription factors known to bind to DNA with GATA4 cooperatively via physical interactions ^27,28^. AlphaFold 3 predicted a heterodimer formation between GATA4 and MEF2C (not shown), implying a potential direct interaction between the two proteins. As the *Gata4* uORF regulates GATA4 protein levels, it is possible that uORF inactivation could impact the GATA4-MEF2 binding dynamics via increasing GATA4 protein expression and altering the stoichiometric ratio of GATA4 to MEF2, thereby modifying enhancer accessibility. This hypothesis gains additional support when considering instances where the identified enhancers act as GATA4 binding sites. Comprehensive biochemical investigations are warranted to enhance our understanding of the GATA4 uORF’s effect on chromatin accessibility.

The physiological and pathological functions of GATA4 have been extensively studied through several *in vivo* models for both reduction and increase of GATA4. Near complete knockout (KO, 70-95% reduction) of Gata4 in the cardiomyocytes resulted in reduced CM hypertrophy, left ventricle (LV) dilation, and reduced ejection fraction (EF) ^7^. Global heterozygous knockout of Gata4 (∼50% permanent reduction) showed mild cardiac dysfunction and sensitivity to TAC-induced pressure overload. A short-term 50% reduction in GATA4 protein expression via a uORF-targeting antisense oligonucleotide (ASO) after TAC surgery reduced CM hypertrophy, prevented CM apoptosis, and improvedbgmn EF^4^. A CM-specific constitutive Gata4 transgene (TG) overexpression model (2.5-fold increase) exhibited late-onset cardiac hypertrophy, fibrosis, reduced fractional shortening (FS), and cardiomyopathy ^6^. An inducible CM-specific Gata4 transgene overexpression (4.6-fold increase) showed mild cardiac hypertrophy, no fibrosis, unaltered FS, and increased angiogenesis ^32^. In sum, the reduction of GATA4 causes eccentric hypertrophy phenotypes. In contrast, overexpression of GATA4 may result in cardiac phenotypes towards concentric hypertrophy (**Figure 6**). Either an increase or a reduction of GATA4 expression in a permanent manner leads to CM hypertrophy, as seen in the present work. However, there is variation in the severity of the cardiac phenotype dependent on the level of reduction or increase in GATA4 as well as the timing and cell specificity of the models. In this work, we generated a relatively mild, global increase in GATA4 expression at the protein translation level (not at the transcription or mRNA level). We have shown a ∼2.2-fold increase in GATA4 protein expression via global inactivation of the GATA4 uORF, causing CM hypertrophy without fibrosis but with increased sensitivity to TAC-induced pressure overload stress. The lack of fibrosis in this global model is interesting, as previous CM-specific overexpression of GATA4 has resulted in a cardiac fibrosis phenotype, which may be driven by constitutive transgenic expression at both transcription and translation levels ^6^. We also note altered enhancer availability after altering only GATA4 protein translation without altering mRNA transcription. Understanding the post-transcriptional regulatory mechanisms of GATA4 expression on enhancer availability and activity may provide novel therapeutic targets in various cardiovascular pathologies.

**Figure 6.**
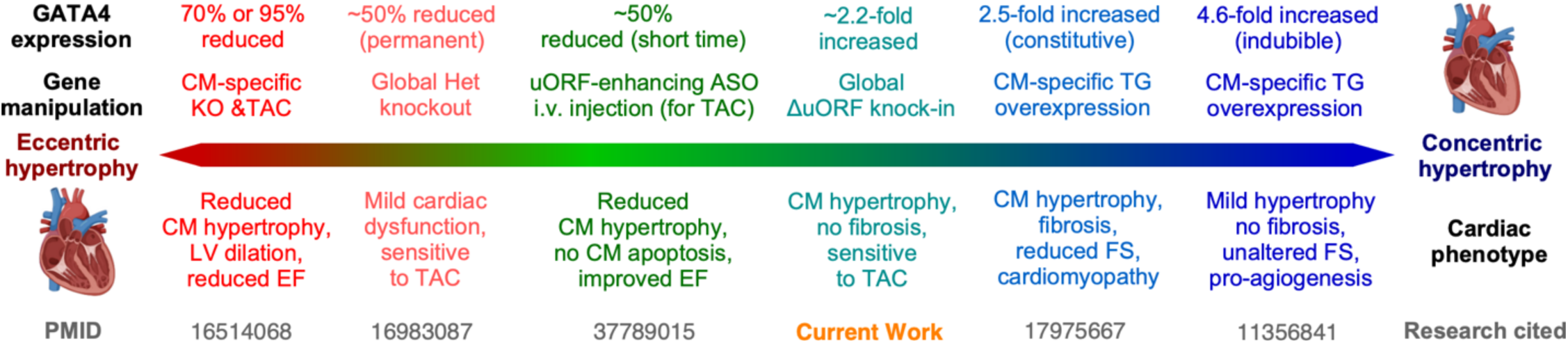
Summary of dose-dependent, GATA4-mediated cardiac physiological and pathological effects following various genetic manipulations. CM: cardiomyocyte; KO: knockout; TAC: transverse aortic constriction; LV: left ventricle; EF: ejection fraction; uORF: upstream open reading frame; ASO: antisense oligonucleotide; i.v.: intravenous; TG: transgenic; FS: fractional shortening.

## Acknowledgments

We appreciate Jiangbin Wu, EngSoon Khor, Jared Hollinger, and Matthew Auguste’s technical assistance in primary mouse adult cardiomyocyte isolation, animal maintenance, and breeding. We appreciate the technical assistance from Erika Flores Medina in histology and Deanne Mickelsen in surgical operations (Aab CVRI). This work was supported in part by the National Institutes of Health (R01 HL132899, HL147954, HL164584, and HL169432 to P.Y.), American Heart Association Established Investigator Award 24EIA1255341 (to P.Y.), Harold S. Geneen Charitable Trust Award from the Medical Foundation at Health Resources in Action (to P.Y.), Rubens Discovery Award 2021 from Aab Cardiovascular Research Institute of University of Rochester Medical Center (to P.Y.), NIH T32 Fellowship (T32 GM068411 to O.M.H.), and National Institutes of Health (R01HL154318 and HL162259 to C.Y.).

## Author contributions

PY launched the study, obtained the funding, and led the project. PY and OMH conceived the ideas, designed the experiments, analyzed the data, and wrote the manuscript. OMH, FJ, UB, AI, YK, SC, and JLS conducted the experimental work. FJ performed the bioinformatic analysis. OMH wrote the initial draft of this manuscript. UB, AI, JLS, YK, and PY revised and finalized the manuscript and worked on the paper revisions. FJ and CY provided technical assistance and conceptual feedback. All the authors discussed the results and had the opportunity to comment on the manuscript.

## Supplementary information

**Supplementary Figure 1.**
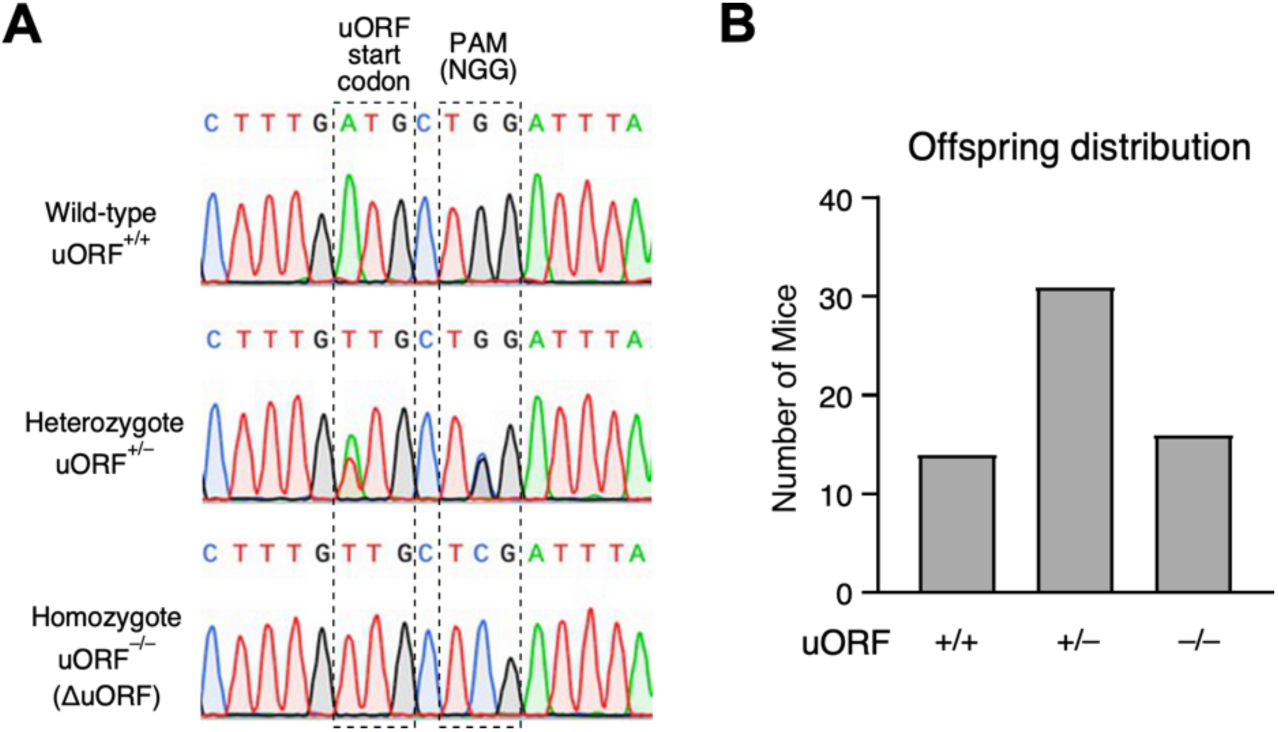
Generation and characterization of *Gata4* ΔuORF mice. **A** Electropherogram displaying DNA sequences obtained from toe clippings of mice with wild-type, heterozygous, and homozygous genotypes, visualized in SnapGene software. **B.** Mendelian distribution of offspring (N = 61; 14 +/+, 31 +/–, 16 –/–) resulting from the breeding of heterozygous mice adheres to the expected 1: 2: 1 ratio of wild-type, heterozygous, and homozygous mice.

**Supplementary Figure 2.**
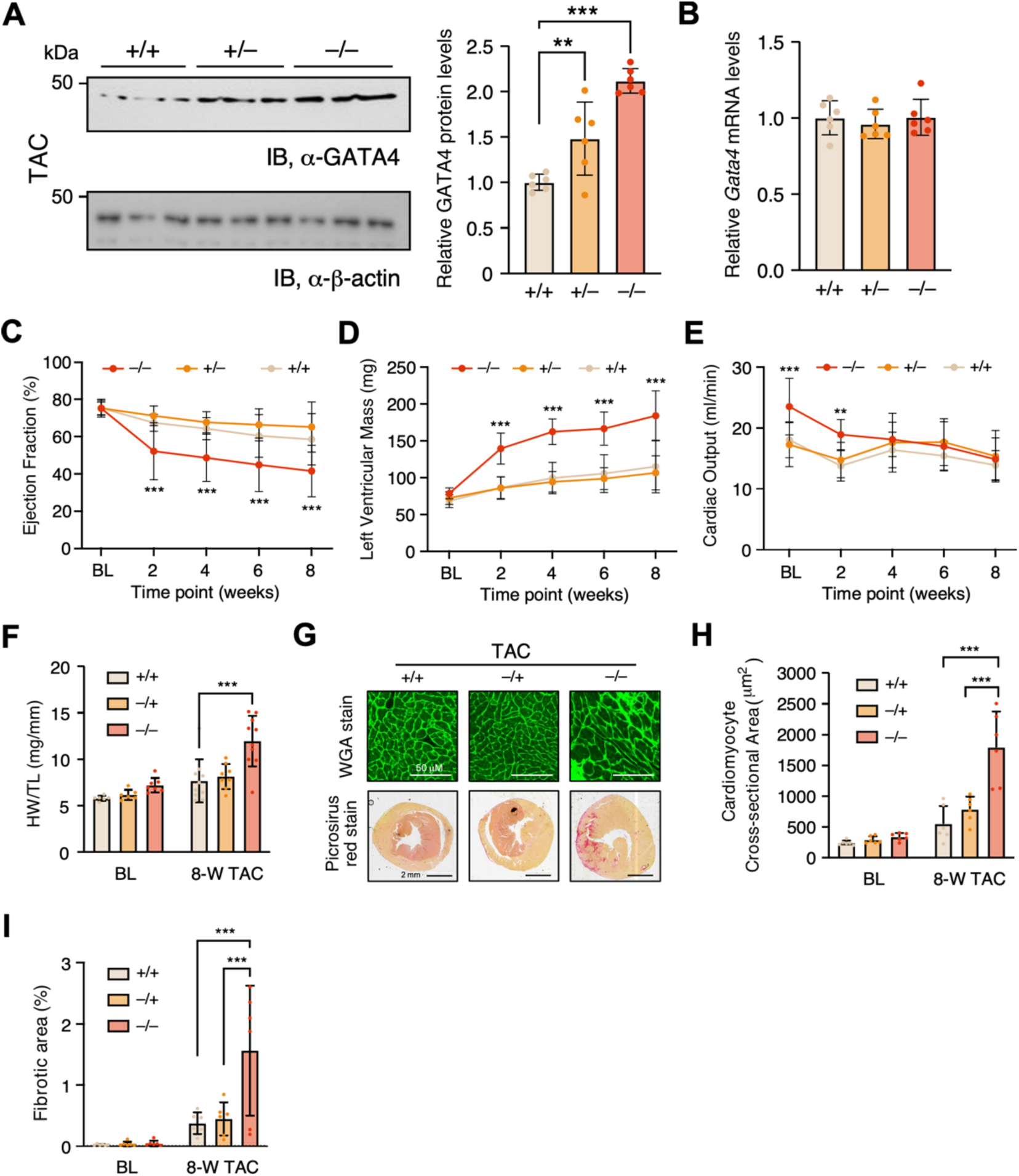
Gata4 ΔuORF mice show increased protein levels and lead to cardiac decompensation in response to transverse aortic constriction surgery. **A.** Western blotting showing the protein expression of GATA4 in wild-type (WT), heterozygous, and homozygous heart lysates normalized to β-actin followed by quantification. **B**. RT-qPCR of *Gata4* mRNA normalized by *Actb* for samples in **A**. **C-E**. Echocardiography parameters over 8 weeks for the mice subjected to TAC surgery. **F-I**. characterizing hypertrophy and fibrosis of the isolated hearts at the 8-week endpoint. **F**. Heart weight normalized to the tibia length as an indicator of hypertrophy. **G**. Upper: Immunomicrograph of mouse cross-sections stained with wheat germ agglutinin (WGA)-Alexa fluor 480 to highlight cellular cross-sectional area, quantified in **H**. Lower: Scan of heart tissue slides stained with picrosirius red, which highlights collagen deposition, quantified in **I**. Data are represented as mean ± SD. ** *P* < 0.05, ** *P* < 0.01, *** *P* < 0.001; Statistical significance was confirmed by 2-way ANOVA followed by Holm-Sidak post hoc test.

**Supplementary Figure 3.**
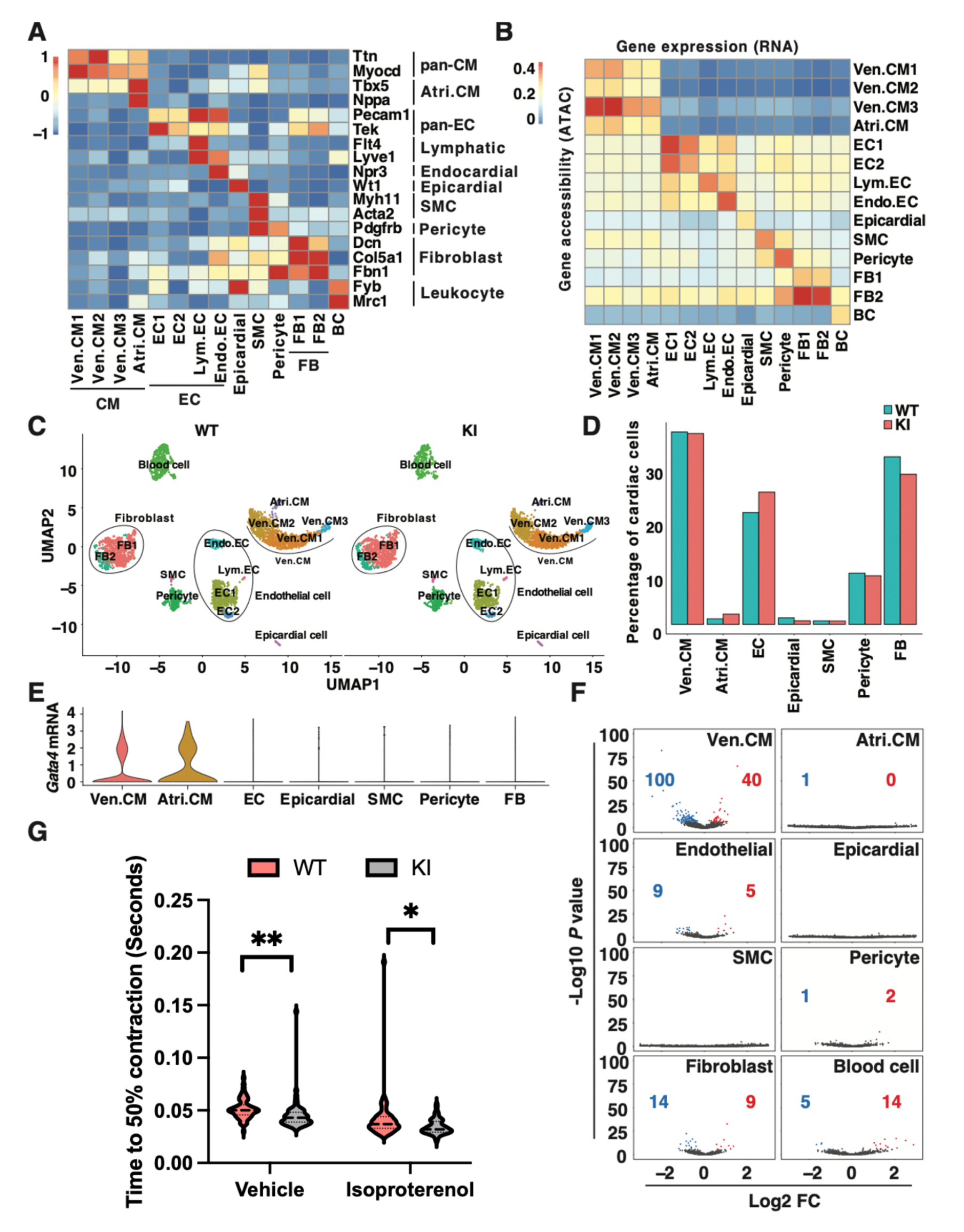
snRNA/ATAC-seq analysis of wildtype and ΔuORF KI mouse hearts. **A.** A heatmap showing the gene accessibilities (summarized chromatin accessibility of promoter and gene body) of marker genes as shown in the Violin Plots in Figure 4C. Scaled and normalized accessibility were plotted. **B**. A heatmap showing the correlation between gene expressions and gene accessibilities within and among clusters. Only the top 3000 variant genes were used to calculate the correlation. **C**. A UMAP presentation of the clustering of wildtype and ΔuORF KI nucleus. **D**. A bar plot showing the percentage of each cardiac cell type in total cardiac cells in wildtype and ΔuORF KI hearts. Blood cells were excluded from the calculation. **E**. Violin Plots showing the *GATA4* mRNA expression in cardiac cells. Scaled and normalized expressions were plotted as the y-axis. **F**. Volcano plots showing the gene expression changes in ΔuORF hearts in each cell type. Genes with significantly increased expression in ΔuORF hearts (Log_2_ fold change > 0.2 and Bonferroni correction adjusted *P*-value < 0.05) were red-colored. Genes with significantly decreased expression (Log_2_ fold change < −0.2 and Bonferroni correction adjusted *P*-value < 0.05) in ΔuORF KI hearts were blue-colored. **G**. A bar graph represents the time cardiomyocytes need to reach 50% of their maximum contraction at baseline (without ISO treatment) and upon isoproterenol (ISO; 10 μM) stimulation.. Ven.CM, ventricular cardiomyocytes; Atri.CM, atrial cardiomyocytes; FB, fibroblasts; EC, endothelial cells; SMC, smooth muscle cells; BC, blood cells.

**Supplementary Figure 4.**
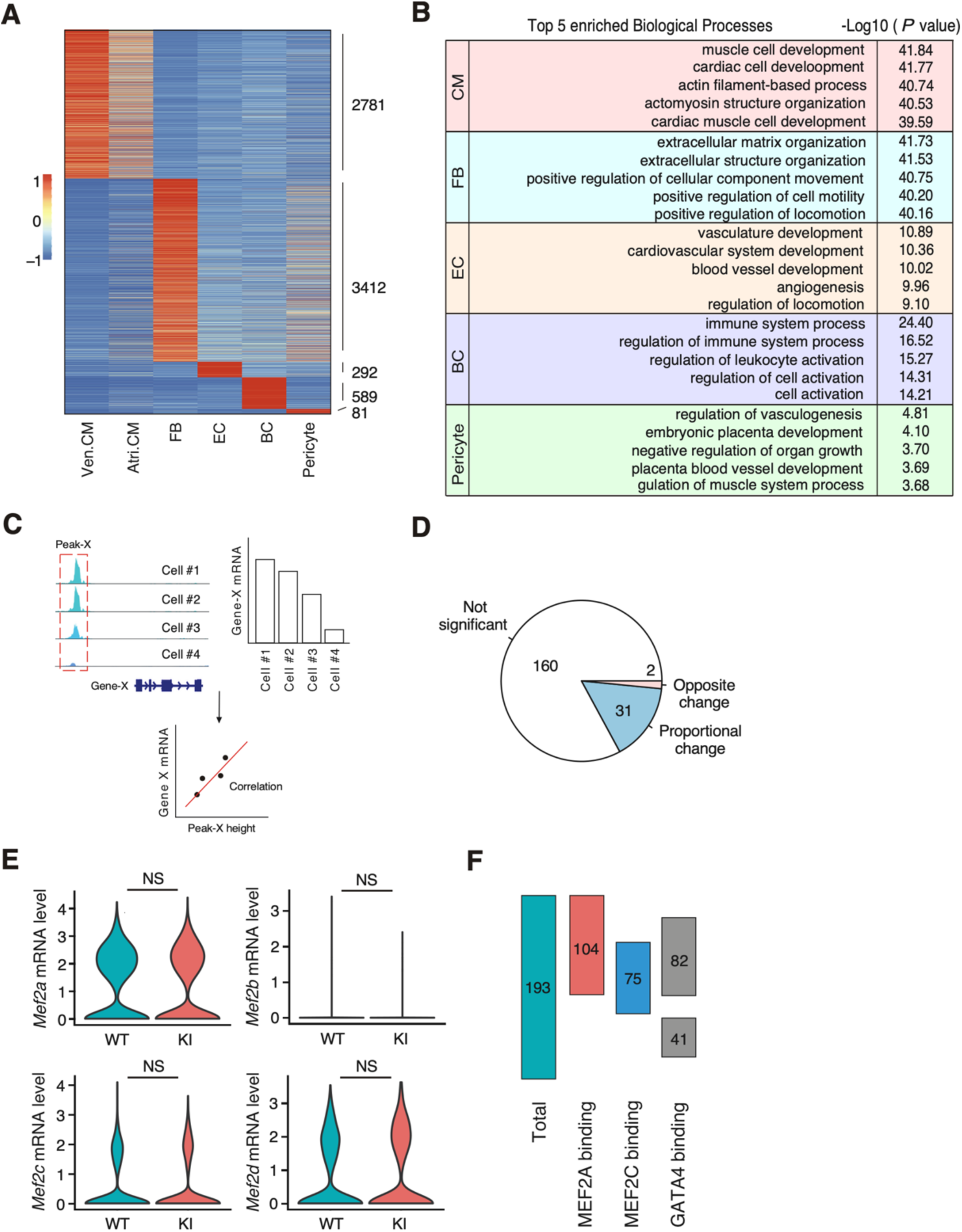
Identification of cis-regulatory DNA elements with altered accessibility in ΔuORF ventricular cardiomyocytes. **A.** A heatmap showing the accessible chromatin regions specific to CM, FB, EC, pericytes, and BC. Scaled and normalized accessibility were plotted. **B**. Genomic Regions Enrichment of Annotations Tool (GREAT) analysis results showing the top 5 enriched biological processes of the cell type-specific regions in Figure S4A. **C**. A sketch showing the workflow of identifying region-gene correlated pairs. The expected coefficient values were calculated for each pair to compute a z-score and *P*-value. Pairs with *P* < 0.05 were considered as region-gene correlated pairs. The Chromatin region in those pairs is treated as pCREs of the correlated genes. **D**. A pie chart shows that gene changes correlated with significantly changed pCREs. 160 of the pCREs with significantly changed accessibility in ΔuORF Ven.CM correlated with genes that have not changed significantly (not significant). 31 of the pCREs change in the same direction as their correlated genes (both increase or decrease in ΔuORF Ven.CM) (coordinate change). 2 of the pCREs change in the opposite direction as their correlated genes (opposite change). **E**. Violin Plots showing the *Mef2a, Mef2b, Mef2c, Mef2d* mRNA expression in Ven.CM. Scaled and normalized expressions were plotted on the y-axis. **F**. A sketch showing the overlap of the significantly changed pCREs and published ChIP-seq data. 104 of the 193 pCREs overlap with MEF2A binding sites, 75 of the 193 pCREs overlap with MEF2C binding sites, and 123 of the 193 pCREs overlap with MEF2A binding sites. CM, cardiomyocytes; FB, fibroblasts; EC, endothelial cells; BC, blood cells.

**Table S1**. Summary of snRNA-seq and snATAC-seq for *Gata4* uORF-inactivating homozygous knock-in (KI) and WT control hearts.

**Table S2**. Marker genes of 14 cell sub-type clusters in snRNA-seq & snATAC-seq.

**Table S3**. Differentially expressed genes (DEGs) in different cell subtypes based on snRNA-seq.

**Table S4**. Cell type-specific peaks for snATAC-seq of KI versus WT hearts.

**Table S5**. Correlations between chromatin regions and mRNA expression of nearby genes.

**Table S6**. Putative cis-regulatory elements (pCREs) with altered accessibility in homozygous ΔuORF KI Ven.CM.

**Table S7**. Overlap regions between snATAC-seq peaks in KI hearts and ChIP-seq identified GATA4 and MEF2 target binding sites in mouse hearts.

## Materials and Methods

### ΔuORF mouse model generation

Single-stranded oligodeoxynucleotides (ssODNs) were engineered to include specific mutations in both the uORF start codon and the PAM sequence to serve as templates for homology-directed repair. The ssODNs were designed with flanking sequences surrounding the targeted editing site. Additionally, guide RNAs (gRNAs) were custom-designed in our lab. This genome-editing strategy was implemented by the University of Auguste Transgenic Mouse Facilities to generate genetically modified mice.

gRNA sequence:

5’-AGAGGTTTCTGCTTTGATGC TGG −3’

ssODN sequence (desired mutations highlighted in lowercase):

5’-ACGATTTCTGGAGCAACCGCAAATCCAATTTGGGATTTTCTTTTTCCTGAGCAAA CCAGAGCCTAGAGGTTTCTGCTTTGtTGCTcGATTTAATTCGTATATATTTTGAGCGAG TTGGGCCTCTCCTCGTTTTTTGATCTCCGGTTG −3’

### Mouse transverse aortic constriction (TAC) cardiac hypertrophy model

For the TAC surgery, mice were anesthetized through continuous administration of inhaled isoflurane (2%) while the surgery was performed. The animals were placed in a supine position, and a midline cervical incision was made to expose the trachea for direct intubation with a 22-gauge plastic catheter. The catheter was connected to a volume-cycled ventilator supplying supportive oxygen. A right thoracotomy was performed. Stenosis was induced using a 27-gauge needle placed on the ascending aorta. Sham-operated mice underwent all aspects of the surgery except for the actual aortic ligature. A ligature was made around the needle and the aorta, completely occluding the aorta. The needle was then removed, causing severe aortic stenosis. Heart rate was monitored during the echocardiography measurement. The stenotic gradient pressure (>100 mmHg) was calculated to evaluate the efficacy of TAC. For the intervention model of TAC surgery in conditional knockout mice, TAC surgery was performed, and a tamoxifen diet was provided for another 6 weeks, starting 14 days post-TAC surgery after a single initial dose of tamoxifen at 30 μg/g of mouse body weight via intraperitoneal injection.

The mice are randomized for experiments. Animal operations, including TAC surgery and echocardiography measurement, were performed blindly by the Microsurgical Core surgeons. Sections and histology analysis were conducted using the Histology Core. For group size justification, we performed a power analysis for One-way ANOVA. As an exemplary experiment, the standard deviation for weight after TAC treatment is about 10%. The minimum difference to be considered significant is 25% in TAC-induced cardiac hypertrophy and heart weight gain. With an overall type I error rate (alpha level) of 5%, at least 5 mice per group are required to achieve 90% power to detect the difference in heart weight. In previous experiences from our Microsurgical Core, we observed a survival rate of ∼85-90% after the TAC procedure. To offset the possible loss of about 1-2 mice per treatment, we used at least 6-7 mice per group.

### Echocardiography

For the TAC surgery mouse model, M-mode short-axis echocardiographic image collection was performed using a Vevo2100 echocardiography machine (VisualSonics, Toronto, Canada) and a linear-array 40 MHz transducer (MS-550D). Heart rate was monitored during echocardiography measurement. Image capture was performed in mice under general isoflurane anesthesia with the heart rate maintained at around 550-650 beats/min. The HR could vary in individual mice due to the potential effect of anesthesia or the surgeon’s operation variation. LV systolic and diastolic measurements were captured from the parasternal short axis in M-mode. Fraction shortening (FS) was assessed as follows: % FS = (end diastolic diameter - end systolic diameter) / (end diastolic diameter) x 100%. Left ventricular ejection fraction (EF) was measured and averaged in both the parasternal short axis (M-Mode) using the tracing of the end diastolic dimension (EDD) and end systolic dimension (ESD) in the parasternal long axis: % EF = (EDD-ESD) / EDD. Hearts were harvested at multiple endpoints depending on the study. In addition to EF and FS, left ventricular end-diastolic diameter (LVEDD), left ventricular end-systolic diameter (LVESD), and wall thickness of left ventricular anterior (LVAWT) and posterior (LVPWT) were also assessed.

### Adult cardiomyocyte isolation

Adult cardiomyocytes (CMs) were isolated from 10-12-week-old male and female mice using a Langendorff perfusion system as previously described ^33^. Mice were fully anesthetized via intraperitoneal injection of ketamine/xylazine. Once the pedal reflex was lost, the mouse was secured in a supine position. The heart was excised and fastened onto the CM perfusion apparatus, and perfusion was initiated in the Langendorff mode. Our Langendorff perfusion and digestion consisted of three steps at 37°C: 4 min with perfusion buffer (0.6 mM KH_2_PO_4_, 0.6 mM Na_2_HPO_4_, 10 mM HEPES, 14.7 mM KCl, 1.2 mM MgSO_4_, 120.3 mM NaCl, 4.6 mM NaHCO_3_, 30 mM taurine, 5.5 mM glucose, and 10 mM 2,3-butanedione monoxime), then switched to digestion buffer (300 U/ml collagenase II [Worthington] in perfusion buffer) for 3 min, and finally perfused with digestion buffer supplemented with 40 μM CaCl_2_ for 8 min. After perfusion, the ventricle was placed in a sterile 35 mm dish with 2.5 ml digestion buffer and shredded into several pieces with forceps. 5 ml stopping buffer (10% FBS, 12.5 μM CaCl_2_ in perfusion buffer) was added and pipetted several times until the tissue dispersed readily, and the solution turned cloudy. The cell solution was passed through a 100 μm strainer. CMs were settled by incubating the cell suspension at 37°C for 30 min. The CMs were resuspended in 10 ml stopping buffer and subjected to several steps of calcium ramping: 100 μM CaCl_2_, 2 min; 500 μM CaCl_2_, 4 min; 1.4 mM CaCl_2_, 7 min. Then the CMs were seeded onto a glass-bottom dish (Nest Biotechnology) pre-coated with laminin (ThermoFisher Scientific). Plates were centrifuged for 5 min at 1,000 g at 4°C to increase adherence, cultured at 37°C for ∼1 hour, and then switched to adult CM culture medium (MEM [Corning] with 0.2% BSA, 10 mM HEPES, 4 mM NaHCO_3_, 10 mM creatine monohydrate, 1% penicillin/streptomycin, 0.5% insulin-selenium-transferrin, and 10 μM blebbistatin for cell culture and downstream assays.

### Measurement of cardiomyocyte contractility

The mechanical properties of cardiomyocytes were assessed using a SoftEdge MyoCam system (IonOptix Corporation, Milton, MA) ^34^. Cardiomyocytes were placed in a chamber and stimulated with a suprathreshold voltage at a frequency of 0.5 Hz. IonOptix SoftEdge software was used to capture changes in sarcomere length during shortening and relengthening. Cell shortening and relengthening were assessed using the following indices: peak shortening (PS), the amplitude myocytes shortened upon electrical stimulation, indicative of peak ventricular contractility; time-to-50% shortening and relaxation, the duration for myocytes to reach 50% shortening and relaxation, indicative of systolic and diastolic duration; and maximal velocities of shortening and relaxation. Intracellular Ca^2+^ was measured using a dual-excitation, single-emission photomultiplier system (IonOptix). Cardiomyocytes were loaded with Fura 2-AM (2 μM) and were exposed to light emitted by an LED lamp through either a 340-nm or 380-nm filter while being stimulated to contract at a frequency of 0.5 Hz. Fluorescence emissions were then detected.

### Single-nucleus ATAC-seq and RNA-seq

Single-nucleus ATAC-seq (snATAC-seq) and snRNA-seq were carried out by SingulOmics Corporation following their established protocols. Whole hearts were harvested from both wild-type and ΔuORF KI mice at the age of 8 weeks (n = 3) and were fast-frozen in liquid nitrogen. Nuclei were isolated from the frozen mouse heart tissue using the 10x Genomics single nuclei isolation kit, adhering to the manufacturer’s instructions. The snRNA-seq and snATAC-seq libraries were simultaneously constructed using the 10x Genomics Chromium System and the 10x Chromium Multiome kit. Each library was subjected to sequencing on the Illumina NovaSeq 6000 platform, generating approximately 200 million paired-end reads (PE150) per library. snRNA-seq and snATAC-seq reads were demultiplexed and aligned to the mouse reference genome mm10 by 10x Genomics Cell Ranger ARC 2.0.2 ^35^. For snRNA-seq, a count matrix was created by summarizing reads mapped to both exon and intron regions of each gene in each nucleus. For snATAC-seq, fragment files were generated containing all unique, properly paired, and aligned fragments. Each unique fragment was associated with a single-cell barcode. A gene accessibility matrix was generated by summarizing fragments mapped to each gene’s promoter and gene body in each nucleus.

### Dimension reduction, clustering, annotation, and differential expression analysis

Only nuclei with more than 200 snRNA-seq reads and snATAC-seq fragments were included in the analysis. Potential doublets were eliminated using DoubletFinder ^36^ with an estimated doublet rate of 5%. Dimension reduction and clustering primarily relied on the top 3000 variant features in snRNA-seq data. Before these processes, data from WT and ΔuORF hearts were integrated, utilizing SCTransformed ^37^ counts and relevant functions in Seruat v4 ^38^ in an R-4.2.2 environment. Dimension reduction was initiated with a Principal Component Analysis (PCA) test, with a subsequent Uniform Manifold Approximation and Projection (UMAP) test utilizing the first 20 principal components. Clustering was performed via the k-nearest neighbors and Shared Nearest Neighbor (SNN) algorithms ^39^. Cluster annotations were initially established by comparing gene expression profiles to publicly available annotated datasets using SingleR ^40^ with gene expression counts normalized and scaled before these comparisons. Reference datasets from Tabula Muris ^41^ containing snRNA-seq data from mouse heart and aorta were employed for this annotation, and additional refinement was undertaken by considering normalized RNA expression and gene accessibility data for well-established cardiac cell-type-specific markers ^42,43^. To identify differentially expressed genes (DEGs) within each cluster, a logistic regression model, known for its robust performance in previous work ^44^, was applied to the normalized counts, with variations in sequencing depth among the nuclei being treated as latent variables. The DEG analysis included only genes expressed in more than 10% of the cluster. Bonferroni correction was applied to adjust the nominal *P* values; genes with |Log2 fold change| > 0.2 and adjusted *P* < 0.05 were considered significant.

### Identification of putative cis-regulatory elements (pCREs)

Putative cis-regulatory elements (pCREs) were identified using Signac v1.1 ^45^ following the developer’s manual. Briefly, snATAC-seq fragments from individual clusters were pooled for peak calling. Peaks were called separately in WT and ΔuORF KI hearts using MACS2 ^46^, and subsequently, the overlapping peaks were included in downstream analysis. The peak height was determined by counting the number of fragments falling into each peak within each nucleus. To facilitate peak height comparisons across different nuclei, these peak heights were normalized using term frequency-inverse document frequency (TF-IDF) normalization ^47^.

The linkages between peaks and genes were established by calculating the correlation between peak height and the RNA expression of genes spanning a 5 × 10^5^ base pair regions encompassing the peak ^48^. The expected coefficient values for the peak are then used to compute a z-score and *P*-value. Peaks displaying a close correlation (*P*-value < 0.05) with the expression of a gene were identified as pCREs for that gene.

To identify pCREs with altered accessibility, a logistic regression test was conducted on the normalized peak heights, as recommended by Signac ^45^. pCREs exhibiting significant accessibility changes underwent motif analysis using a hypergeometric test in Signac ^45^. The analysis aimed to assess the enrichment of transcription factor binding motifs sourced from JASPAR2023 ^49^. BEDtools ^50^ were employed to perform an overlap analysis of pCREs with published ChIP-seq regions ^26^.

### Molecular Cloning

pCRE sequences were amplified via PCR, using primers designed to incorporate a KpnI cut site in the forward primer and an XhoI cut site in the reverse primer. After amplification, the PCR fragments were purified using a PCR purification kit (ThermoFisher Scientific). These purified fragments were co-digested in the same reaction mixture with the pGL3-SV40 Promoter vector, using KpnI and XhoI restriction enzymes (NEB). One microliter of the digested mixture was subsequently incubated with T4 DNA ligase (NEB) and transformed into competent bacterial cells.

### Dual-luciferase reporter assay

Transfected cells were incubated with Dual-Glo luciferase substrate (Promega) according to the manufacturer’s instructions. The final readings of the FLuc were normalized to RLuc to obtain the relative luminescence reading.

